# Quantitative analysis of the frequency of chromosome loss following DSB induction

**DOI:** 10.1101/2025.01.15.633104

**Authors:** Seiya Matsuno, Ryo Ishida, Ryotaro Kawasumi, Kouji Hirota, Takuya Abe

## Abstract

Numerical abnormalities in chromosomal states, referred to as aneuploidy, is commonly observed in many cancer cells. Although numerous internal and external factors induce aneuploidy, the primary cause of aneuploidy in humans remains unclear. DNA damage is identified as a potential cause of aneuploidy by inducing chromosome segregation errors. However, a direct relationship between DNA damage and aneuploidy remains poorly understood. A major reason for this is the extremely low frequency of aneuploidy in cultured cells, making quantitative analyses challenging. In this study, we investigated the relationship between DNA damage and aneuploidy in cell lines containing minichromosomes. These chromosomes are more prone to loss than normal chromosomes, with the rate of loss substantially increased following exposure to various DNA-damaging agents. To determine whether damaged chromosomes were subjected to direct loss or whether chromosome loss occurred as an indirect consequence of a prolonged G2 phase or other factors, we used the CRISPR-Cas9 system to introduce a single DNA double-strand break (DSB) on a minichromosome. The rate of minichromosome loss increased by approximately seven-fold compared with that of the control. Furthermore, the loss rate was significantly elevated in the absence of KU70, a key factor in non-homologous end joining, and upon inhibition of ataxia telangiectasia mutated (ATM), a DNA damage checkpoint protein. Finally, two closely spaced nicks, believed to generate a 5’-overhang, were also shown to induce minichromosome loss. These findings indicated that a single DSB or two closely spaced nicks can cause aneuploidy if improperly repaired in vertebrates.

## 1. Introduction

Aneuploidy refers to an increase or decrease in the number of chromosomes from the normal state. Considered an important hallmark of cancer, the study of its causes and effects is clinically relevant. Aneuploidy primarily arises from errors during somatic cell division and is often attributed to various factors, including abnormal sister chromatid cohesion, defective spindle checkpoints, increased centrosome numbers, abnormal kinetochore-microtubule attachments, and kinetochore malformations ^1^. In addition, radiation-induced DNA damage can induce aneuploidy ^2^. One mechanism by which DNA damage induces aneuploidy is by increasing the number of centrosomes in damaged cells, resulting in the pulling of chromosomes toward three or more spindle poles ^3^. Another hypothesis suggests that DNA damage affects the microtubules and kinetochores. For instance, DNA damage caused by radiation activates damage checkpoints, stabilizes the binding between kinetochores and microtubules, and leads to errors in chromosomal segregation ^4^. Moreover, factors involved in the damage checkpoint have been proposed to regulate the spindle checkpoint during DNA damage or replication ^5^. Furthermore, DNA damage has been linked to abnormal sister chromatid cohesion ^6^, and the activation of damage checkpoint pathways is thought to influence kinetochore flexibility ^7^. Despite these findings, the various mechanisms underlying aneuploidy resulting from DNA damage remain unclear.. A single DNA double-strand break (DSB) is known to cause chromosome loss in both human and mouse embryos ^8,9^. Chromosome loss following DSB induction by Cas9 has been attributed to the formation of chromosome bridges between daughter nuclei, potentially resulting in incorrect separation of sister chromatids and errors in chromosome distribution ^10^. Studies have indicated chromosome loss as a byproduct of genome editing, but the underlying mechanism and the factors influencing its rate remain poorly understood. To further investigate the relationship between DNA damage and aneuploidy, we utilized an artificially developed minichromosome from chicken DT40 cells. Minichromosomes are ideal tools for quantitative measurements of chromosome loss owing to their inherent instability compared to normal chromosomes. Although various methods for measuring chromosomal aneuploidy have been reported, we developed a novel system to quantify aneuploidy by using flow cytometry to detect the presence or absence of mCherry fluorescence introduced into minichromosomes. This system was employed to assess chromosomal loss induced by DNA damaging agents using the CRISPR-Cas9 system.

### 2. Materials and methods

A complete list of reagents and detailed methodology are available in Supplementary data.

## 3. Result

### 3.1 DNA-damaging agents induce minichromosome loss

Previously, we established a cell line with one of the three copies of chromosome 2s in chicken DT40 cells edited into a minichromosome following sequential transfection with two telomere-seeding vectors ^11^. The minichromosome contained a *GFP* gene, which enables quick measurement of the minichromosome loss rate using flow cytometry. In accordance with the previous notation, the GFP-harboring minichromosome cell line was denoted as DT40 Chr2 (1,1,mini-GFP). We used this cell line to evaluate the effects of DNA damage on chromosomal loss quantitatively. First, the effects of various DNA-damaging agents, including methyl methanesulfonate, cisplatin, etoposide, ICRF193, and camptothecin were examined. The cells were exposed to various concentrations of each drug for 48 h and the number of cells and GFP loss rates were measured simultaneously using flow cytometry. To ensure consistent concentration conditions for these drugs, the concentration at which cell viability was 10% (IC90) was determined by calculating the relative cell number at each concentration. All five DNA-damaging agents exhibited increased minichromosome loss rates at their respective IC90 (Fig. 1A). Thus, the minichromosome system can detect DNA damage-induced chromosomal losses with high sensitivity.

**Fig 1.**
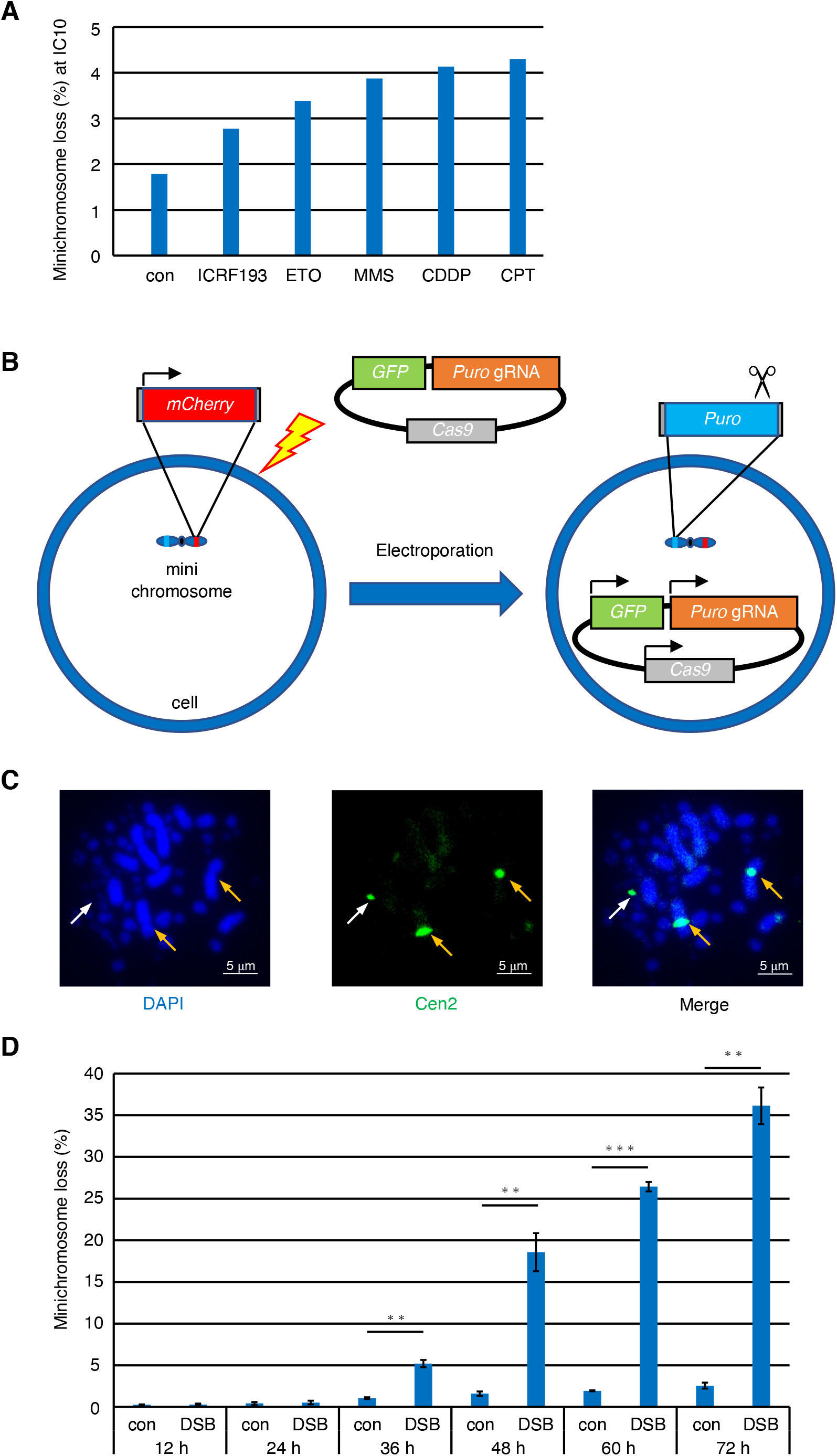
Double strand break on mini-chromosome increases mini-chromosome loss (A) DT40 Chr2 (1, 1, mini-GFP) cells were pre-incubated with puromycin and L-histidinol for 24 h to eliminate cells lacking the minichromosome. Puromycin and L-histidinol were washed out, and the cells were exposed to DNA-damaging agents. Following 2 days of culture, cells were analyzed using flow cytometry. (B) A minichromosome-specific DSB was introduced using the CRISPR-Cas9 system. A gRNA expression vector harboring *cas9* and *gfp* was transfected into cells via electroporation. (C) Representative images showing the presence of the minichromosome by FISH. White arrow represents minichromosome, orange arrow represents chromosome 2. (D) After eliminating cells lacking minichromosome by puromycin and L-histidinol, DT40 Chr2 (1, 1, mini-mCherry) cells were transfected with a gRNA expression vector (designated ‘DSB’) or the control vector without gRNA (designated ‘control’). Cells were then cultured for indicated times, and minichromosome loss rates were analyzed using flow cytometry. Error bars indicate the standard deviation (SD) of the mean from three independent experiments. From left to right, ^**^*P*=0.0047, ^**^*P*=0.0072, ^***^*P*=0.00015, ^**^*P*=0.0010, one-tailed t-test.

### 3.2 A single DSB on the minichromosome induces chromosome loss

Although DNA-damaging agents increase minichromosome loss rates, it remains unclear whether DNA damage directly induces chromosome loss on minichromosomes or whether the loss is indirectly mediated via the DNA damage signaling pathway, for example, through prolonged G2 phase via checkpoint activation, given that DNA damage agents induce DNA lesions randomly throughout the genome. To determine this, we examined the effect of site-specific DNA damage on minichromosomes. CRISPR-Cas9 system is not only widely used as a genome editing tool ^12^ but is also useful for inducing site-specific DNA damage ^13^. To analyze cells expressing only gRNA and *cas9* gene, we generated a cell line expressing *mCherry* on the minichromosome and a gRNA expression vector bearing *cas9* and *gfp* (Fig. 1B). Generation of the DT40 Chr2 (1,1,mini-mCherry) cell line was validated via FISH using a probe targeting the centromeric sequence of chicken chromosome 2 (Fig. 1C). A gRNA was designed based on a puromycin resistance gene (*puro*) and a vector expressing only *cas9* and *gfp* was used as a control. Initially, minichromosome loss rates were measured every 12 h after transfection, and a time-dependent increase was observed (Fig. 1D). As the amount of *cas9*-expressing cells (estimated by GFP expression) decreased gradually 48 h after transfection, minichromosome loss rates were measured at this time point in subsequent experiments.

### 3.3 A DSB-induced minichromosome loss occurs in a cell cycle-dependent manner

Chromosome loss is thought to occur as a result of nuclear division failures, even when triggered by DNA damage. To examine whether DSB-induced minichromosome loss occurs during cell division, we used hydroxyurea (HU) to slow cell cycle progression via inhibition of DNA replication. Compounds that arrest the cell cycle at mid-metaphase, such as microtubule inhibitors, induce minichromosome loss. As shown in Supplementary Fig. 1, the rate of DSB-induced minichromosome loss decreased with increasing concentrations of HU, suggesting that minichromosome loss occurred in a cell cycle-dependent manner, potentially during cell division.

### 3.4 A DSB on a different chromosome does not induce minichromosome loss

Although we showed that DSB on a minichromosome induced minichromosome loss, it is possible that activated checkpoints or other DNA damage response pathways indirectly induced minichromosome loss. To elucidate this, we introduced DSB on a different chromosomes. To employ the same gRNA designed to target *puro*, we generated a cell line in which *puro* was replaced by a blasticidin S resistance gene (*bs*), and *puro* was targeted to an *OVA* locus on the p-arm of chromosome 2 (Supplementary Fig. 2A). Induction of a DSB on the p-arm of chromosome 2 did not increase the minichromosome loss rate, suggesting that minichromosome loss is directly induced by DSB on the minichromosomes (Fig. 2A).

**Fig 2.**
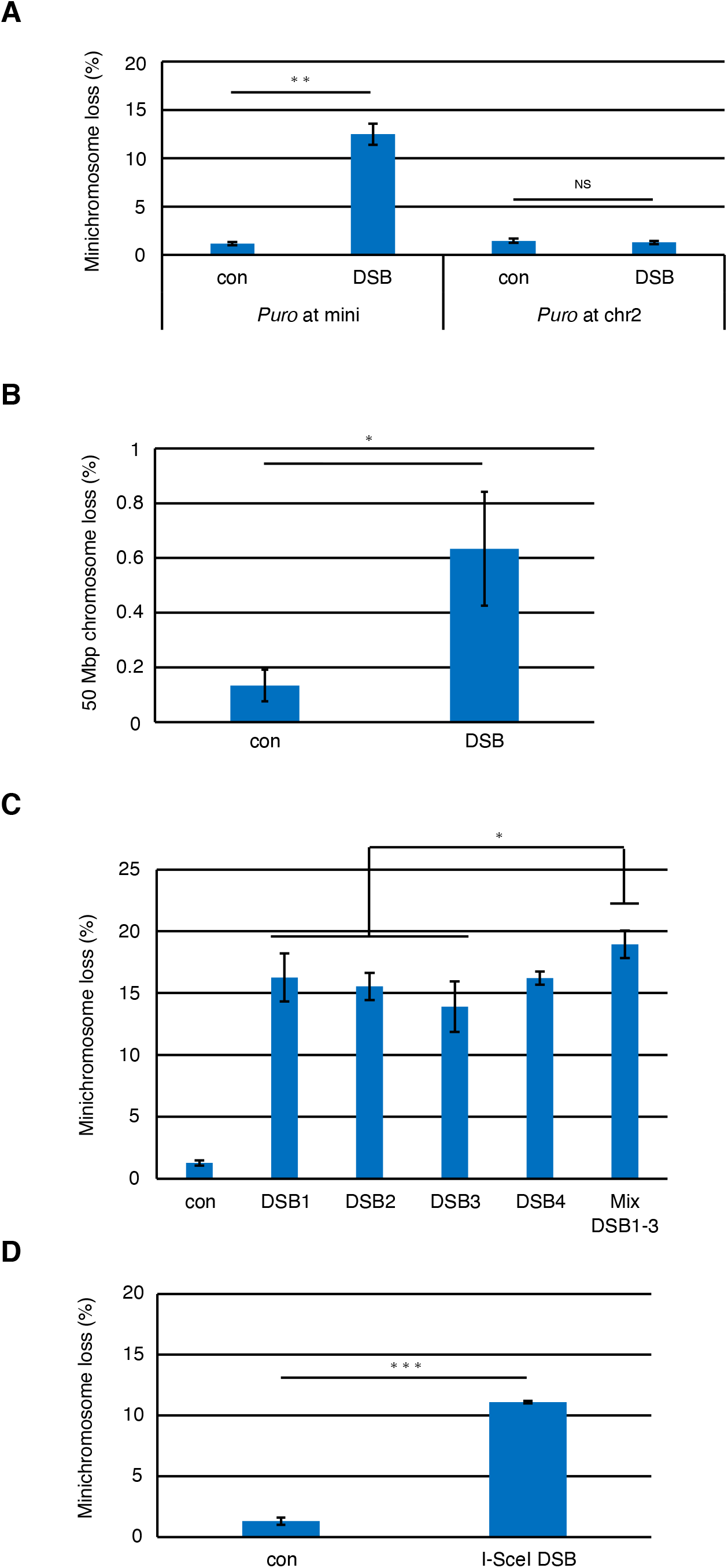
DSB directly induces chromosome loss. (A) A DSB was induced on minichromosome (*Puro* at mini) or full-length chromosome 2 (*Puro* at chr2). ^**^*P*=0.0026, one-tailed t-test. (B) Chromosome loss rates were measured as in Fig. 1D for DT40 Chr2 (1,1,50 Mb-mCherry) cells. ^*^*P*=0.038, one-tailed t-test. (C) Mini-chromosome loss rates were measured as in Fig. 1D for DT40 Chr2 (1, 1, mini-mCherry) cells with each of the four gRNA represented in Supplementary Fig. 3A or the mixture of DSB1-3. The gRNA DSB1 is used in other experiments. From left to right,^*^*P*=0.032, ^*^*P*=0.023, ^*^*P*=0.030, one-tailed t-test. (D) DT40 Chr2 (1, 1, mini-mCherry) cells harboring an I-SceI recognition site on mini-chromosome was transfected with an *I-SceI* expression vector or a control vector. Minichromosome loss rates were analyzed using flow cytometry. ^***^*P*=0.00045, one-tailed t-test.

Next, we generated a cell line, DT40 Chr2 (1,1,50 Mb-mCherry), where one of the three chromosome 2s used for minichromosomes generation was edited to 50 Mb in size to examine whether a DSB was capable of inducing chromosome loss in normal-size chromosomes (Fig. 2B and Supplementary Fig. 2B). Despite a much lower chromosome loss rate compared to minichromosomes, DSB-induced chromosome loss occurred in the 50 Mb chromosome. This result indicates that DSB-induced chromosome loss is not specific to minichromosomes.

### 3.5 An increase in DSBs further elevates the minichromosome loss rates

To exclude the possibility that minichromosome loss was induced specifically by the gRNA used in Fig. 1D, we designed three additional gRNAs for *puro* (DSB2-4) (Supplementary Fig. 3A). As shown in Fig. 2C, all four gRNAs designed based on *puro* induced a comparable degree of minichromosome loss. Using the gRNAs, we next examined whether an increase in DSB numbers further boosts minichromosome loss rates. Notably, increasing the dosage of single gRNA plasmid did not affect the minichromosome loss rates (Supplementary Fig. 3B). Although the increase in minichromosome loss was not additive when a mixture of three gRNA’s mixture (DSB1-3) was used, the increase rates were statistically significant compared to those when using a single gRNA (Fig. 2C). Therefore, multiple DSBs occurring at nearby sites may increase the risk of chromosome loss.

### 3.6 An I-SceI-mediated DSB induces minichromosome loss

Although the above results show that DSB induced by Cas9 increases the minichromosome loss rate, this effect may be specific to Cas9, which generates a blunt-ended DSB facilitated by a gRNA. To examine whether the phenomenon is generalized to all forms of DSBs, we employed another nuclease, I-SceI, a homing enzyme that recognizes an 18 bp specific sequence and generates a 3’ overhang DSB ^14^. We generated a cell line with an I-SceI recognition sequence adjacent to *puro* on a minichromosome. *I-SceI* was expressed by transfection of the *I-SceI* expression vector, and minichromosome loss rates were measured. I-SceI-induced DSB increased the minichromosome loss rate (Fig. 2D). Given that the expression level and cutting efficiency differ between I-SceI and Cas9, comparing the efficiency of minichromosome loss between them is challenging. However, the results clearly showed that DSB-induced minichromosome loss was not specific to Cas9.

### 3.7 NHEJ-deficiency increases minichromosome loss after a DSB induction

Following DSB induction, DNA damage responses, including checkpoint and DNA repair pathways, are activated. Cells have two mechanisms for DSB repair: non-homologous end joining (NHEJ) and homologous recombination (HR)^15^. During NHEJ, cut ends of DNA are recognized by KU70/80 and DNA-PKcs, processed by Artemis, and ligated by the LIG4/XRCC4/XLF complex in a template-independent manner ^16^. NHEJ is an error-prone DNA repair pathway, whereas HR uses an intact DNA template to repair the break and restore the original sequence ^17^. To assess the minichromosome loss rates in the absence of these DNA repair pathways, we generated *KU70*- and *XRCC3*-deficient cells harboring minichromosomes. Although the rate of spontaneous chromosome loss was not affected, DSB-induced chromosome loss increased in *KU70*-deficient cells (Fig. 3A). In contrast, the rate of DSB-induced chromosomal loss was decreased in *XRCC3*-deficient cells (Fig. 3B). Thus, we can conclude that NHEJ suppresses chromosome loss by immediately ligating the broken ends, whereas HR induces segregation errors, potentially through the generation of an HR-dependent chromosome bridge ^18^.

**Fig 3.**
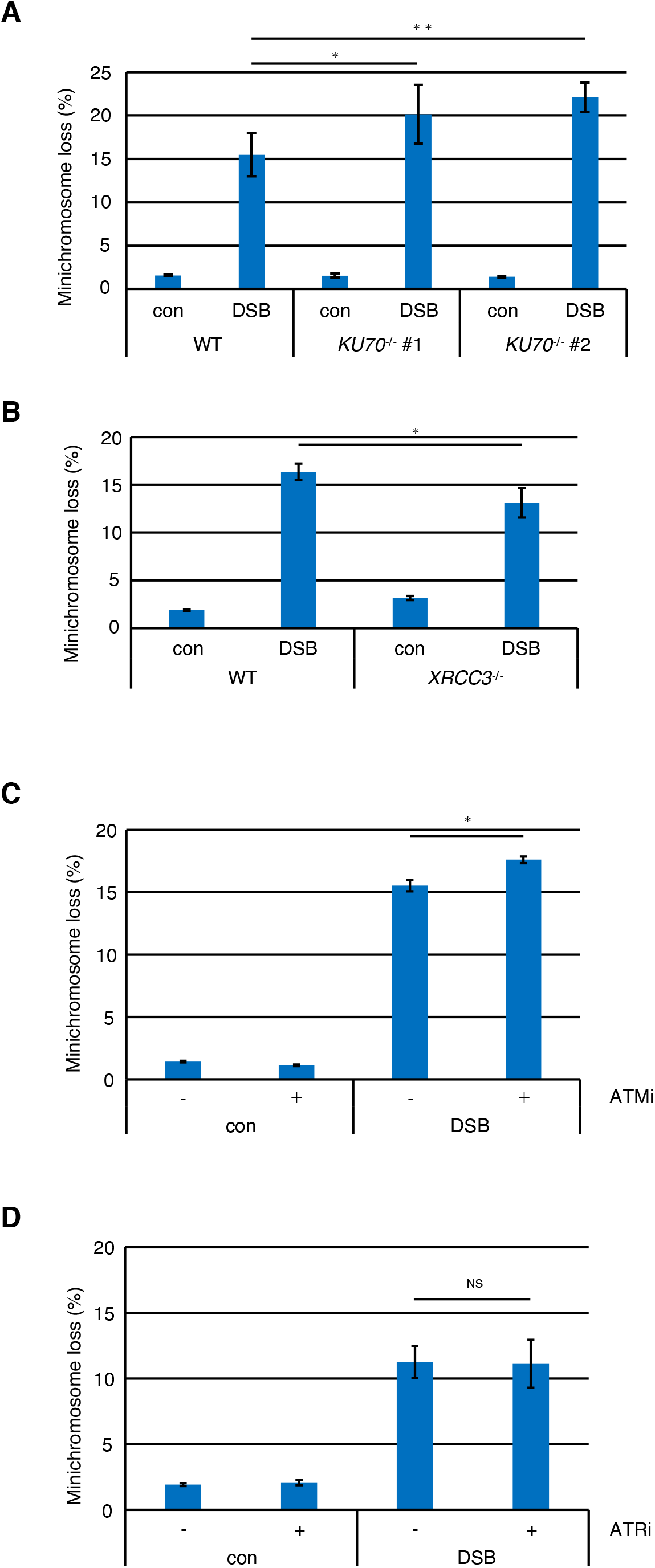
Inhibition of NHEJ or ATM increases DSB-induced mini-chromosome loss. (A) A DSB was introduced on *puro* as in Fig. 1D, and mini-chromosome loss rates were compared between DT40 Chr2 (1,1,mini-mCherry) (shown as ‘WT’) and *KU70*^*-*/*-*^ DT40 Chr2 (1,1,mini-mCherry) (shown as ‘*KU70*^*-*/*-*^’) backgrounds. Error bars indicate the standard deviation (SD) of the mean from four independent experiments. ^*^*P*=0.047, ^**^*P*=0.0011, one-tailed t-test. (B) Minichromosome loss rates were compared between ‘WT’ and *XRCC3*^*-*/*-*^ DT40 Chr2 (1,1,mini-mCherry) (shown as ‘*XRCC3*^*-*/*-*^’) backgrounds. ^*^ *P*=0.017, one-tailed t-test. (C-D) Mini-chromosome loss rates were measured as in Fig. 1D for DT40 Chr2 (1, 1, mini-mCherry) cells untreated or treated with KU60019 (C) ^*^*P*=0.036, one-tailed t-test. or VE-821 (D)

### 3.8 Checkpoint-deficiency increase minichromosome loss following DSB induction

Chromosome segregation is regulated by different checkpoint proteins during cell division to ensure accurate separation of sister chromatids ^19^. DSBs activate two major checkpoint kinases, ATM and ATR, to prevent cell cycle progression ^20^. To examine the contribution of the DNA damage checkpoint in preventing chromosome loss in our system, we used ATM (KU-60019) and ATR (VE-821) inhibitors. Although neither inhibitor affected the rate of spontaneous minichromosome loss, KU-60019 significantly increased the rate of DSB-induced minichromosome loss (Fig. 3C). In contrast, VE-821 had no effect (Fig. 3D). This suggests that ATM-dependent checkpoint activation prevents minichromosomal loss following DSB induction.

### 3.9 Mps1 inhibition and DSB induction further induce minichromosome loss

The spindle assembly checkpoint (SAC) is a critical safeguard during cell division that prevents aneuploidy. Mps1 is a conserved kinase that regulates SAC by phosphorylating SAC proteins ^21^ and those involved in mitosis, including Mad1, Cdc31, and Spc98^22^. To examine whether SAC prevents chromosome loss, we used the Mps1 inhibitor, BAY1217389^23^. Initially, we inoculated cells with various concentrations of BAY1217389 and found it to induce minichromosome loss (Supplementary Fig. 4A). Because BAY1217389 is highly cytotoxic at concentrations > 12 nM, we used this concentration in subsequent experiments. The combination of Cas9-induced DSB and BAY1217389 exposure resulted in an additional increase in chromosome loss (Supplementary Fig. 4B), indicating that SAC does not protect against DSB-induced chromosome loss.

### 3.10 Cas9 pair nickase increases minichromosome loss

Cas9 contains two nuclease domains, HNH and RuvC, which cleave DNA strands complementary and non-complementary to the 20 nucleotide guide sequence in crRNAs, respectively ^24^. We assessed minichromosome loss using an RuvC-inactivating variant (Cas9 D10A) that generates DNA nicks instead of DSBs. The same four gRNA sequences depicted in Supplementary Fig. 3A were used to generate the DNA nicks. Although slight increases were observed in several gRNAs (Nick2, Nick3 and Nick4), the increase was substantially smaller than that induced by DSBs. Genome editing by double nicks using a pair of gRNAs specific for the complementary strand minimizes off-target mutagenesis ^25^, and thus considered a safer genome editing tool compared to wild-type Cas9 using a single gRNA ^26^. Using Nick2 as the reference point, the effect of pair nicking on minichromosome loss was measured, combined with a nick on the same strand side (Nick3), and two nicks on opposite strand (Nick1 and Nick4). The combination of Nick1 and Nick2 and Nick2 and Nick3 induced minichromosome loss comparable to Nick2 alone; however, the combination of Nick2 and Nick4 induced a higher level of minichromosome loss (Fig. 4A). Genome editing efficiency was substantially different between protospacer-adjacent motif (PAM)-out designs, where the PAM sequences were located at both ends of the target region, and PAM-in designs, where PAM sequences were positioned closer to the center of the target region (Supplementary Fig. 5)^25^. To further confirm whether the PAM-out designs increased minichromosome loss, we designed another gRNA (Nick5) (Fig. 4B) and examined four PAM-out designs. All four combinations induced approximately 10% minichromosome loss (Fig. 4C). These results suggest that paired nicking in PAM-out designs efficiently induces minichromosome loss.

**Fig 4.**
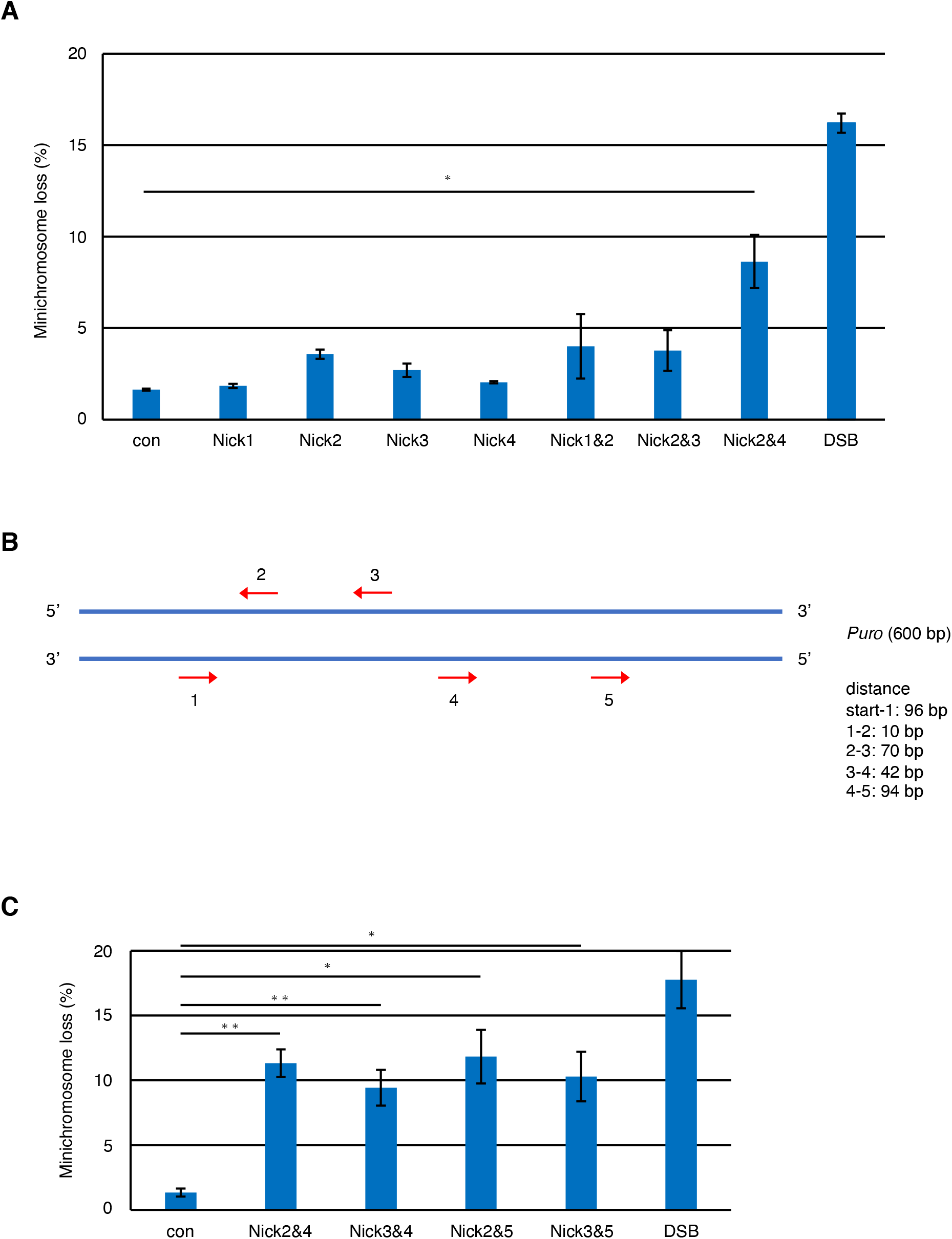
Mini-chromosome loss is induced by dual nicks. (A) Mini-chromosome loss caused by single or double nicks. Mini-chromosome loss rates were measured as in Fig. 1D for DT40 Chr2 (1, 1, mini-mCherry) cells with Cas9 or Cas9 D10A and indicated numbers of gRNA. The numbers of gRNA correspond to those in Supplementary Fig. 3A. ^*^*P*=0.015, one-tailed t-test. (B) The direction and location of five gRNAs designed based on *puro*. (C) Minichromosome loss rates were measured as in Fig. 4A for DT40 Chr2 (1, 1, mini-mCherry) cells with Cas9 or Cas9D10A and indicated combinations of gRNA. From left to right, ^**^*P*=0.0057, ^**^*P*=0.0099, ^*^*P*=0.012, ^*^*P*=0.015, one-tailed t-test.

## 4 Discussion

In this study, we employed a sensitive minichromosome system and demonstrated that a DSB-generated Cas9 or two close nicks in a specific direction by Cas9 D10A induces chromosome loss. Additionally, DSB-induced chromosome losses were observed when I-SceI was used in place of Cas9 and a 50 Mb chromosome was used instead of a minichromosome, indicating that the phenomenon is not limited to Cas9 or minichromosomes. Moreover, the deficiency of NHEJ, but not HR, resulted in an increase in DSB-induced minichromosome loss. The non-requirement of HR in the system can be attributed to Cas9 cutting both strands of sister DNA after replication, leaving no available template DNA for HR. Thus, it remains unclear whether HR plays a pivotal role in preventing chromosome loss in DSBs in natural conditions. A deficiency in NHEJ causes a minor increase in the rate of minichromosome loss (∼20%), likely because other repair pathways compensated for this deficiency. Polθ is a potential repair system candidate that compensates for the NHEJ deficiency. The alternative end-joining (Al-EJ) mediated by Polθ mediates mitotic DSB repair ^27^. Considering that the activity of NHEJ is suppressed during mitosis ^28^ and that Cas9 induces DSBs even during mitosis ^29^, Polθ-mediated mitotic DSB repair may play a crucial role in preventing chromosome loss.

Our results showed that Cas9-mediated minichromosome loss was further enhanced by ATM inhibition. The ATM-mediated G2 checkpoint plays a crucial role in preventing cell cycle progression to mitosis until the DNA repair process is complete. Moreover, a previous study using the I-SceI-glucocorticoid receptor (GR) fusion protein demonstrated that a DSB is sufficient to activate the checkpoint and that ATM influences DSB-induced sister chromatid cohesion during metaphase ^30^. Integrating these findings, the checkpoint may prevent DSB-induced chromosome loss by slowing metaphase progression via the accumulation of cohesin as well as by inducing cell cycle arrest at the G2 phase.

Nickase-mediated double nicks are safer and more precise genome editing system than Cas9, because it minimizes off-target mutagenesis. However, the effect of these two nicks on chromosomal loss has not been thoroughly investigated. Based on our findings, we propose that two nicks in close proximity, as designed in the PAM-out direction, efficiently induce chromosome loss. In the PAM-out direction, nicks were positioned outside the two Cas9 molecules, whereas in the PAM-in direction, nicks were sandwiched between the two Cas9 molecules (Supplementary Fig. 5). The duration for which Cas9 D10A remains bound to the DNA strand after the induction of nicks remains uncertain. However, if Cas9 remains bound for an extended period, it could potentially inhibit the binding of DNA repair factors. Considering that nicks designed in the PAM-out direction result in higher genome editing efficiency, they may provide better access for various DNA repair factors, including nucleases, potentially resulting in the conversion of the two nicks into a DSB. Further studies are required to elucidate the detailed mechanisms underlying the high frequency of chromosomal loss in the PAM-out direction.

## Glossary

PAM-in design: PAM sequences positioned closer to the center of the target region
PAM-out design: PAM sequences located at both ends of the target region

## Acknowledgement

We thank Rika Rifana Sari and M. Nakagawa for technical assistance; Editage (www.editage.jp) for English language editing.

## Author contributions: CRediT

Conceived and designed the experiments: T.A. Data analysis: S.M., R.K. and K.H. Performed the experiments: S.M., R.I. and T.A. Wrote the paper: S.M and T.A.

## Funding

This work was supported by Grants from the Uehara Memorial Foundation, the Mochida Memorial Foundation for Medical and Pharmaceutical Research, Senri Life Science Foundation, A-STEP from JST (JPMJTM22BQ) and JSPS KAKENHI (20K06760 and 22H05072) to TA.

## Data Statement

The data that support the findings of this study are available from the corresponding author, T.A., upon reasonable request.

## Notes

### Competing Interest Statement

The authors have declared no competing interest.

## References

1. Wilhelm, T., Said, M. & Naim, V. DNA Replication Stress and Chromosomal Instability: Dangerous Liaisons. Genes (Basel) 11, 642 (2020).

2. Eastmond, D. A. & Pinkel, D. Detection of aneuploidy and aneuploidy-inducing agents in human lymphocytes using fluorescence in situ hybridization with chromosome-specific DNA probes. Mutation Research/Environmental Mutagenesis and Related Subjects 234, 303–318 (1990).

3. Dodson, H. et al. Centrosome amplification induced by DNA damage occurs during a prolonged G2 phase and involves ATM. EMBO J 23, 3864–3873 (2004).

4. Bakhoum, S. F., Kabeche, L., Murnane, J. P., Zaki, B. I. & Compton, D. A. DNA-Damage Response during Mitosis Induces Whole-Chromosome Missegregation. Cancer Discov 4, 1281–1289 (2014).

5. Kim, E. M. & Burke, D. J. DNA Damage Activates the SAC in an ATM/ATR-Dependent Manner, Independently of the Kinetochore. PLoS Genet 4, e1000015 (2008).

6. Masamsetti, V. P. et al. Replication stress induces mitotic death through parallel pathways regulated by WAPL and telomere deprotection. Nat Commun 10, 4224 (2019).

7. Strecker, J. et al. DNA damage signalling targets the kinetochore to promote chromatin mobility. Nat Cell Biol 18, 281–90 (2016).

8. Zuccaro, M. V. et al. Allele-Specific Chromosome Removal after Cas9 Cleavage in Human Embryos. Cell 183, 1650-1664.e15 (2020).

9. Papathanasiou, S. et al. Whole chromosome loss and genomic instability in mouse embryos after CRISPR-Cas9 genome editing. Nat Commun 12, 5855 (2021).

10. Leibowitz, M. L. et al. Chromothripsis as an on-target consequence of CRISPR-Cas9 genome editing. Nat Genet 53, 895–905 (2021).

11. Abe, T., Suzuki, Y., Ikeya, T. & Hirota, K. Targeting chromosome trisomy for chromosome editing. Sci Rep 11, 18054 (2021).

12. Cong, L. et al. Multiplex Genome Engineering Using CRISPR/Cas Systems. Science (1979) 339, 819–823 (2013).

13. Liu, M. et al. Global detection of DNA repair outcomes induced by CRISPR-Cas9. Nucleic Acids Res 49, 8732–8742 (2021).

14. Jacquier, A. & Dujon, B. An intron-encoded protein is active in a gene conversion process that spreads an intron into a mitochondrial gene. Cell 41, 383–394 (1985).

15. Scully, R., Panday, A., Elango, R. & Willis, N. A. DNA double-strand break repair-pathway choice in somatic mammalian cells. Nat Rev Mol Cell Biol 20, 698–714 (2019).

16. Lieber, M. R. The Mechanism of Human Nonhomologous DNA End Joining. Journal of Biological Chemistry 283, 1–5 (2008).

17. Thompson, L. H. & Schild, D. Homologous recombinational repair of DNA ensures mammalian chromosome stability. Mutation Research/Fundamental and Molecular Mechanisms of Mutagenesis 477, 131–153 (2001).

18. Chan, Y. W., Fugger, K. & West, S. C. Unresolved recombination intermediates lead to ultra-fine anaphase bridges, chromosome breaks and aberrations. Nat Cell Biol 20, 92–103 (2018).

19. Barnum, K. J. & O’Connell, M. J. Cell Cycle Regulation by Checkpoints. in 29-40 (2014). doi:10.1007/978-1-4939-0888-2_2.

20. Smith, H. L., Southgate, H., Tweddle, D. A. & Curtin, N. J. DNA damage checkpoint kinases in cancer. Expert Rev Mol Med 22, e2 (2020).

21. Chmielewska, A. E., Tang, N. H. & Toda, T. The hairpin region of Ndc80 is important for the kinetochore recruitment of Mph1/MPS1 in fission yeast. Cell Cycle 15, 740–747 (2016).

22. Liu, X. & Winey, M. The MPS1 Family of Protein Kinases. Annu Rev Biochem 81, 561–585 (2012).

23. Wengner, A. M. et al. Novel Mps1 Kinase Inhibitors with Potent Antitumor Activity. Mol Cancer Ther 15, 583–592 (2016).

24. Gasiunas, G., Barrangou, R., Horvath, P. & Siksnys, V. Cas9-crRNA ribonucleoprotein complex mediates specific DNA cleavage for adaptive immunity in bacteria. Proceedings of the National Academy of Sciences 109, (2012).

25. Ran, F. A. et al. Double Nicking by RNA-Guided CRISPR Cas9 for Enhanced Genome Editing Specificity. Cell 154, 1380–1389 (2013).

26. Nakajima, K. et al. Precise and efficient nucleotide substitution near genomic nick via noncanonical homology-directed repair. Genome Res 28, 223–230 (2018).

27. Gelot, C. et al. Polθ is phosphorylated by PLK1 to repair double-strand breaks in mitosis. Nature 621, 415–422 (2023).

28. Orthwein, A. et al. Mitosis Inhibits DNA Double-Strand Break Repair to Guard Against Telomere Fusions. Science 344, 189–193 (2014).

29. Rodriguez-Muñoz, M., Serrat, M., Soler, D., Genescà, A. & Anglada, T. Breakage of CRISPR/Cas9-Induced Chromosome Bridges in Mitotic Cells. Front Cell Dev Biol 9, (2021).

30. Dodson, H. & Morrison, C. G. Increased sister chromatid cohesion and DNA damage response factor localization at an enzyme-induced DNA double-strand break in vertebrate cells. Nucleic Acids Res 37, 6054–6063 (2009).

