## Supplementary Figure for "Quantitative analysis of the frequency of chromosome loss following DSB induction"

### Supplementary Figure 1

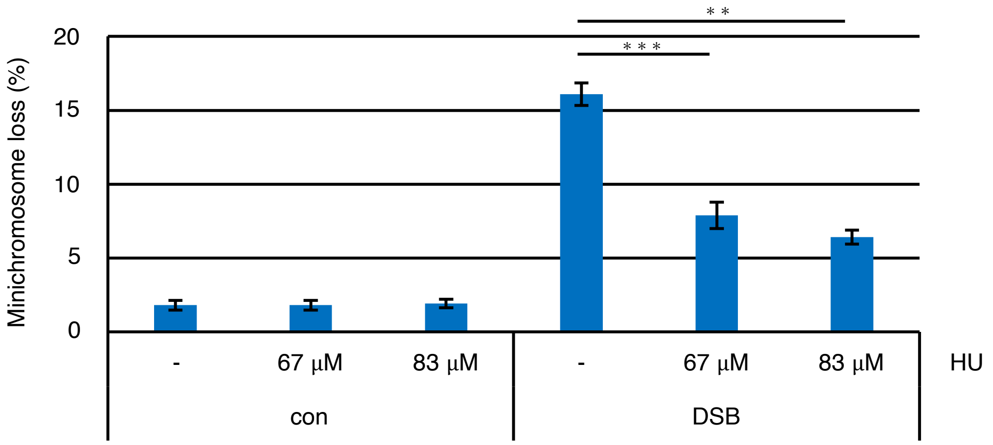

### Supplementary Figure 2

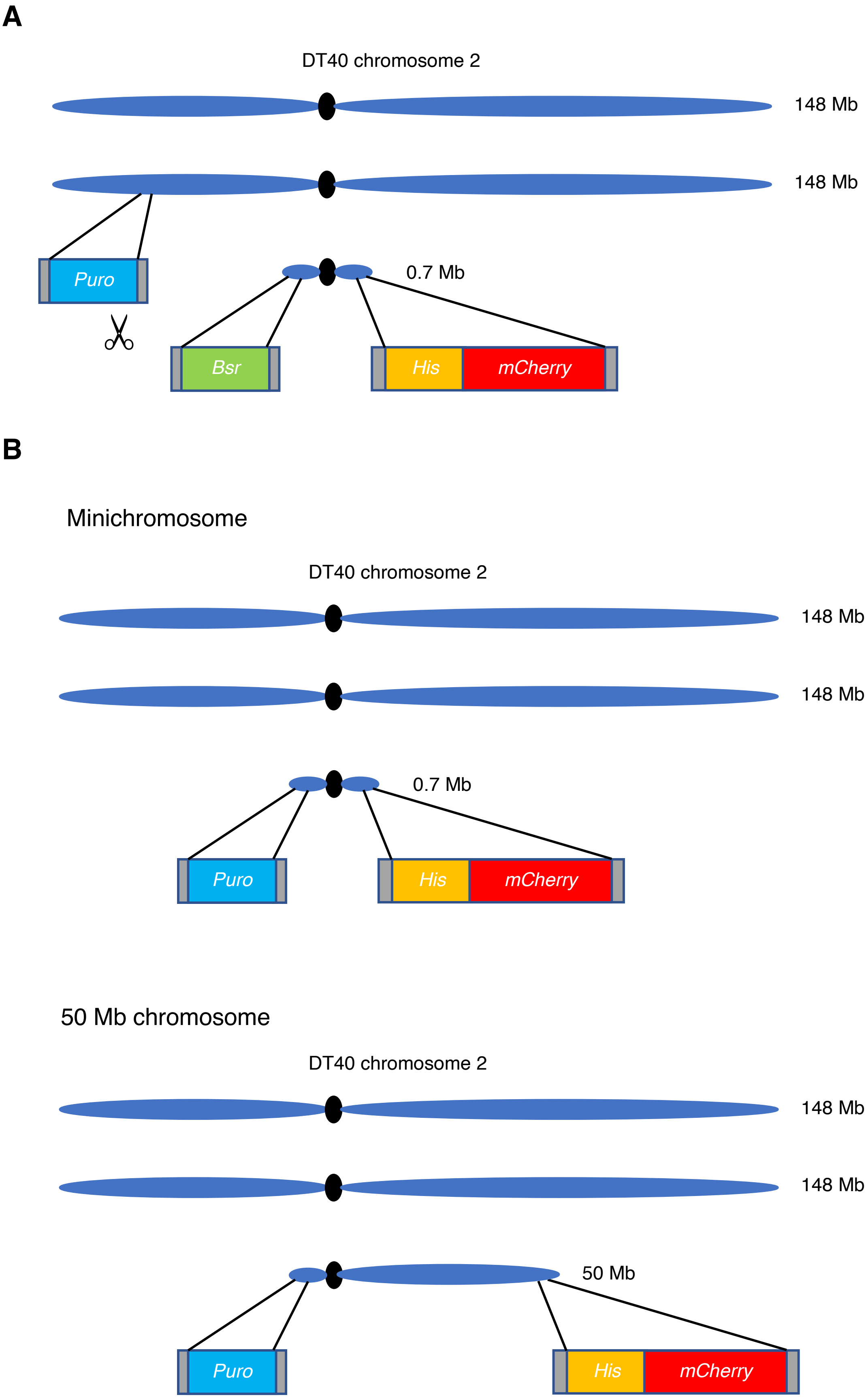

### Supplementary Figure 3

**A**

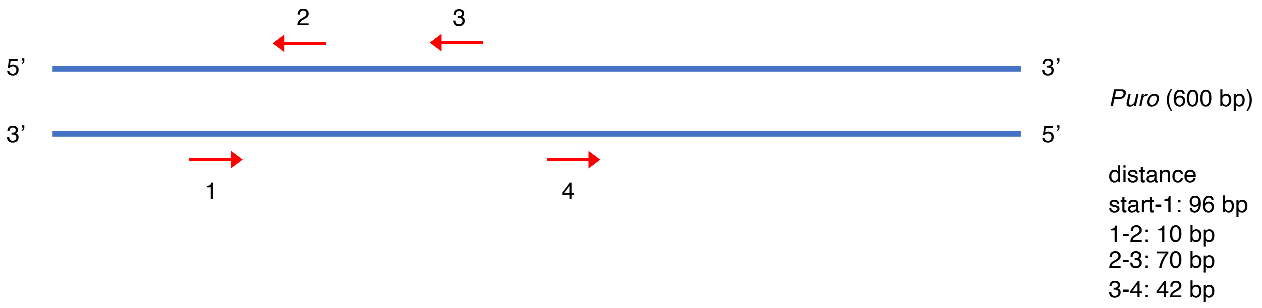


**B**

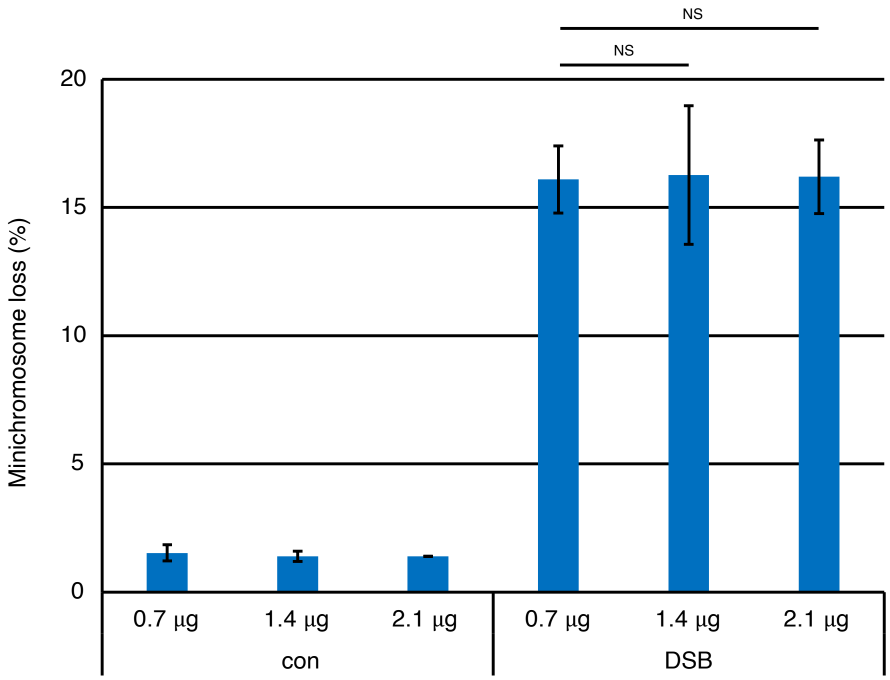

### Supplementary Figure 4

**A**

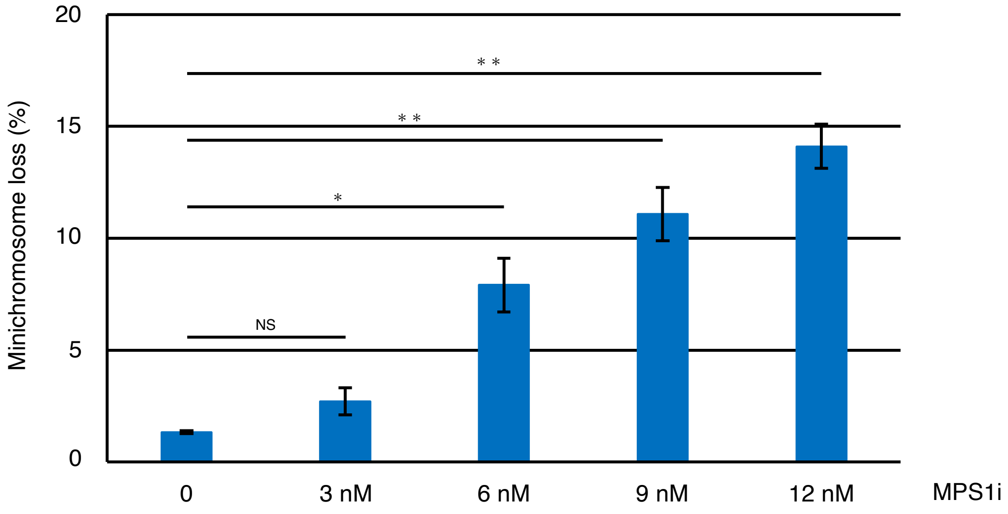


**B**

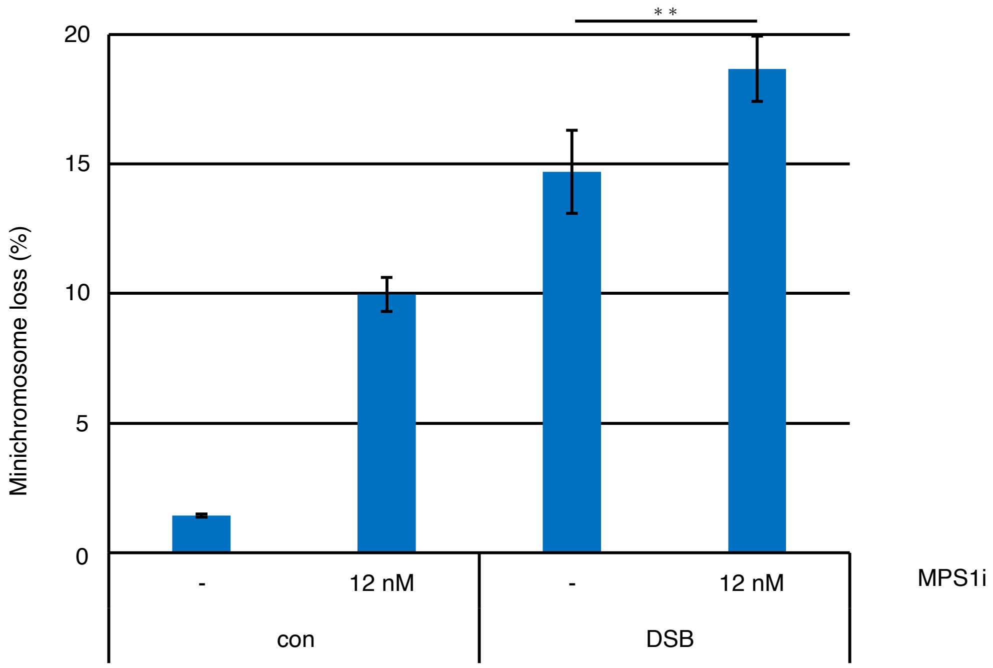

### Supplementary Figure 5

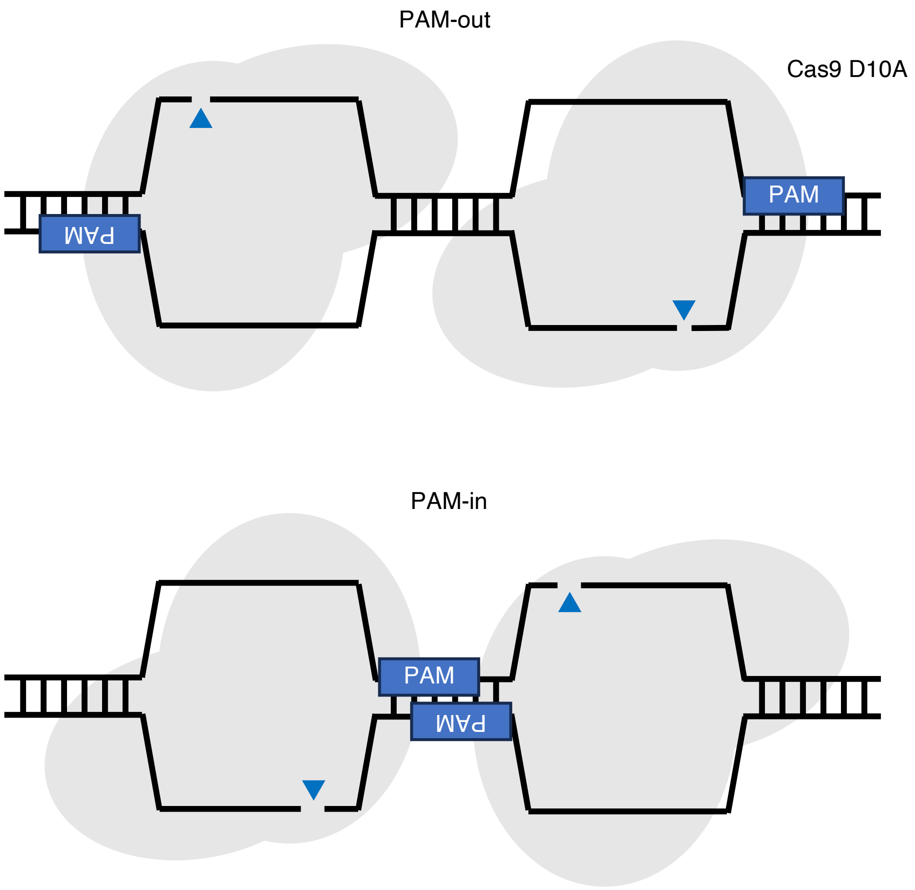

### Supplementary Table 1

| Name | Sequence |
|---|---|
| Puro1 | CACGCGCCACACCGTCGACCCGG |
| Puro2 | GCTCGGTGACCCGCTCGATGTGG |
| Puro3 | CGTGGTCCAGACCGCCACCGCGG |
| Puro4 | CCGCGCATGGCCGAGTTGAGCGG |
| Puro5 | TCGCCCGACCACCAGGGCAAGGG |

#### Materials and methods

##### Cell culture

DT40 cells were cultured at 39.5ºC in Dulbecco's modified Eagle’s medium F12 (Wako) supplemented with 10% fetal bovine serum (Biowest), 2% chicken serum (Gibco), penicillin/streptomycin mix (Nakalai), 2 mM L-glutamine (Nakalai) and 10 µM 2-mercaptoethanol.

##### Plasmid construction and transfection

The pX330 vector (Addgene plasmid #42230) [1]was used to express the CRISPR-Cas9 system. The gRNA sequences are listed in Supplementary Table 1. KU70 KO vectors [2], an OVA KI vector [3], and telomere seeding vectors targeting the *EGFR* and *TPK1* loci were used to generate minichromosome-harboring cells, as previously described [4]. These vectors were subsequently transfected into DT40 cells using the Neon® Transfection System (Invitrogen) according to the manufacturer’s instructions.

*I-SceI* site knock-in vector was generated by modifying the *EGFR* telomere-seeding vector [4]. Two DNA oligos, 5’-CTAGTtagggataacagggtaatA-3’ and 5’-CTAGTattaccctgttatccctaA-3’ containing an *I-SceI* recognition site were annealed and cloned into the SpeI site of the *EGFR* telomere seeding vector.

The telomere seeding vector was used to generate a 50 Mb chromosome from chromosome 2, incorporating genomic PCR products combined with a hisD selection marker cassette. The homology arm was amplified using the primers 5′-aaaGCGGCCGCcctaactaacaaccctctgctaaatc-3′ (NotI) and 5′-aaaACTAGTgcgcacacctgctactgacaagg-3′ (SpeI). The amplified PCR products were cloned into a telomere repeat-harboring vector [4].

##### Fluorescence in situ hybridization (FISH)

The probes were prepared by PCR amplification of the centromeric repeat sequence of chromosome 2 from chicken DT40 cells and labeled with Green 496 dUTP according to the Nick Translation DNA Labeling System 2.0 (Enzo Life Sciences, Inc.). Cells were subjected to 0.1 µg/mL colcemid treatment for 2 h, to increase the number of metaphase-arrested cells. These cells were collected and treated with 75 mM KCl for 13 min at room temperature and then fixed in a methanol-acetic acid solution (3:1) for 30 min. Cell suspensions were dropped onto ice-cold glass slides and stored at -30°C. Chromosome plates were sequentially transferred into containers filled with 2× saline-sodium citrate (SSC), 4% formaldehyde, water, and ethanol, and then dried at 37°C. Chromosomal DNA was denatured with a denaturing solution at 75°C. After washing with ethanol and acetone, the preparations were dried at 37°C. Labeled probes were denatured, mixed with hybridization buffer, dropped onto the slides, and hybridized overnight at 37°C. The hybridized slides were washed with 2× SSC at 42°C, followed by incubation with 1× phosphate buffered saline. Finally, 2 µg/mL DAPI was added dropwise. Images were captured using a BZ-X810 fluorescence microscope (Keyence, Tokyo, Japan).

**Reference**


1. Cong, L. *et al.* Multiplex Genome Engineering Using CRISPR/Cas Systems. *Science (1979)* **339**, 819–823 (2013).
2. Takata, M. *et al.* Homologous recombination and non-homologous end-joining pathways of DNA double-strand break repair have overlapping roles in the maintenance of chromosomal integrity in vertebrate cells. *EMBO J* **17**, 5497–5508 (1998).
3. Fukushima, T. *et al.* Genetic Analysis of the DNA-dependent Protein Kinase Reveals an Inhibitory Role of Ku in Late S–G2 Phase DNA Double-strand Break Repair. *Journal of Biological Chemistry* **276**, 44413–44418 (2001).
4. Abe, T., Suzuki, Y., Ikeya, T. & Hirota, K. Targeting chromosome trisomy for chromosome editing. *Sci Rep* **11**, 18054 (2021).
